# Utilization of Cell-penetrating Peptide Adaptors to Enhance Delivery of Variably Charged Protein Cargos

**DOI:** 10.64898/2026.03.09.710683

**Authors:** Daniel P. Morris, Nathaniel I. Turner, Jojo J. Croffie, Jonathan L. McMurry

## Abstract

Cell-penetrating peptides (CPPs) can deliver biomacromolecular cargos into cells, potentially enabling a new mode of intracellular drug delivery. However, a major problem with CPP-mediated delivery is entrapment of CPPs within endosomes and covalent linkages ensure CPPs and cargos share a common fate. We previously developed a CPP-adaptor system based on reversible, calcium-dependent cargo binding that produces cargo release from adaptors as complexes dissociate following internalization and Ca^2+^ efflux from early endosomes. Having employed CPP-adaptors with an array of protein cargos of differing charges, it became apparent that positively charged cargos often appeared to dominate internalization and that association with the adaptor had little effect. To systematically address the effects of cargo charge and CPP function, we tested the ability of several adaptors to increase internalization of a set of adaptor binding GFP cargos having net charges of +9, +15, +20, +25 and +36. Intrinsic internalization of these cargos reproduced reported patterns showing that positive charge increases internalization. However, labeling these cargos with a chemical fluorophore revealed that GFP fluorescence grossly underestimated total internalization. Internalization was charge and concentration dependent with more positive cargos showing apparent saturation of internalization at 100-400 nM, well below the concentrations at which covalently linked CPP-cargos are dosed. We tested the ability of 5 adaptors to internalize these cargos. Our prototype adaptor, TAT-CaM, was completely ineffective with the +9 cargo, but internalized moderately charged cargos extremely efficiently at concentrations far below the µM range. A derivative adaptor, TAT-LAH4-CaM, was highly effective with all cargos and produced similar maximal internalization at 100-400 nM. However, two adaptors specifically designed with increased positive charge inhibited internalization of the most positive cargos. One of these, GFP-CaM, based on the supercharged GFP with net charge of +36, did increase internalization of the least positive cargos, demonstrating an adaptor with high affinity for the cell surface can increase internalization of a neutral cargo at very low concentration. The common maximal level of intrinsic GFP cargo internalization correlated with surface loading of these cargos, suggesting a limit to the beneficial effects of increased plasma membrane association. However, TAT-CaM further increased internalization via an apparently distinct mechanism. In this limited study of the interaction of cargo charge and adaptor efficacy, we found diverse behaviors that hint at the power and flexibility possible with adaptor/cargo internalization.

## Introduction

Inability to cross membranes is a common failure of otherwise promising therapeutic leads. Cell-penetrating peptides (CPPs) have long held promise for overcoming these failures by efficient, nonviral transmembrane delivery of biomolecular “cargos” to the cytoplasm or other subcellular destination wherein their therapeutic targets are located. Classically, a CPP is covalently attached to a protein or other biomolecule. The CPP binds to cell surface proteoglycan receptors, triggering endocytosis and bringing the cargo along with it [1]. However, it has long been recognized that cargo delivery to the cytoplasm is very inefficient because CPP-cargos get trapped in endosomes and are either recycled to the cell surface or targeted for degradation. This is perhaps the major reason that no CPP-based drug has yet been approved by the U.S. FDA despite dozens of clinical trials [2].

To overcome this endosomal escape problem, we developed CPP-adaptors that spontaneously form noncovalent complexes with cargos via Ca^2+^-dependent coupling. Our prototype, TAT-CaM, consists of the well-characterized CPP TAT [3, 4] N-terminally fused to calmodulin [5]. Other adaptors have different CPPs, different EF hand proteins, and/or additional features such as endosomal escape enhancing peptide (also called endolytic peptide, EP) sequences [6, 7]. CPP-adaptors bind calmodulin binding sequence (CBS)-containing cargo proteins with low nM affinity in the presence of Ca^2+^ and negligibly in its absence [5, 8]. Thus, CPP-adaptor/CBS-cargo complexes bind tightly in extracellular media and during internalization but dissociate after formation of endosomes as Ca^2+^ efflux from endosomes occurs during trafficking [9], leaving cargo to egress to the cytoplasm while the adaptors remain trapped like other CPPs [6, 7, 10, 11]. While dissociation enhanced cargo escape remains a critical adaptor function, our use of CPP-adaptors also suggested that separation of cargo and adaptor function provided a powerful model for investigation of CPP-mediated delivery.

It is well established that positive charge is part of the mechanism used by CPPs to gain cell entry [12]. Evidence suggests this is due to association of positive CPPs with negatively charged proteoglycans on the cell surface [13, 14]. Very recently, an unbiased genetic screen was used to identify target cell proteins that aid internalization of TAT-associated cargos, identified proteins were responsible for synthesis of proteoglycans, providing very strong support for this idea [15]. It has also been recognized that “supercharged” green fluorescent proteins (GFPs) [16, 17], in which surface residues have been altered to increase positive surface charge, act as cell-penetrating proteins (CPPrs) with modest intrinsic internalization [18]. This points to the obvious fact that CPPs fused to cargo proteins display internalization properties clearly dependent on both the CPP and cargo charge, greatly confusing determination of CPP efficacy. The interaction of CPP function and intrinsic cargo internalization due to positive charge has not been systematically investigated.

Because our CPP-adaptors allow separation of intrinsic cargo internalization from the effects of CPP-adaptor/cargo complexes, the interaction of cargo charge and CPP-adaptor function can easily be compared. Based on the supercharged GFPs mentioned above, we created sets of cargos with CBS-GFPs having net charges of 9, 15, 20, 25 and 36. Using confocal microscopy we investigated internalization of these cargos alone and in complex with a set of 5 adaptors previously shown to be efficacious [7].

Two improved adaptors used an increase in net positive charge to mitigate the negative effects on internalization of the strongly negative calmodulin. One of these, TAT-NMR-CaM, was based on *Heterocephalus glaber* calmodulin, which includes an additional positive domain amino terminal to the conserved calmodulin core sequence [10]. The second, GFP-CaM, dispensed with the TAT sequence and used the supercharged GFP with a net +36 as the sole CPPr [7].

The other two adaptors were initially designed to increase endosomal escape by incorporating putative endolytic sequences between TAT and CaM. The LAH4 sequence in TAT-LAH4-CaM dramatically increased internalization of cargos based on maltose binding protein, which has a net negative charge. Although LAH4 was identified as an endolytic peptide and numerous reports have since shown it has CPP functionality [19, 20]. The second adaptor, TAT-AUR-CaM, incorporated the Aurein 1.2 sequence which has endolytic properties in the BHK cell line used here [21]. This adaptor displayed decreased ability to internalize the MBP cargos. Further, a cargo with the Aurein 1.2 sequence in complex with GFP-CaM became trapped in peripheral vesicles [7]. This suggests a mechanism for the endolytic properties of the sequence, which could become potent for TAT-AUR-CaM in complex with strongly internalizing cargo.

Comparing internalization of supercharged GFP cargos with differing net positive charges produced outcomes that were cargo charge- and concentration-dependent. A maximal level of internalization was associated with apparent saturation of surface binding, suggesting the major factor necessary for internalization can be positive charge. On the other hand, TAT-CaM and TAT-LAH4-CaM further increased internalization suggesting an additional mechanism of action.

## Materials & Methods

### Plasmids

Plasmids used were previously described [7] or constructed as described in Supporting Information (S1 Fig). *E. coli-*optimized synthetic genes were designed, synthesized and cloned (GeneScript, Piscataway, NJ, USA) into NdeI or NcoI and BamHI sites in pET19b (EMD Millipore, USA) with an in-frame stop codon prior to the BamHI site. In the case of CPP-adaptors, the N-terminal His tag was followed by the TAT peptide sequence (YGRKKRRQRRR). GFP-CaM consists of sequence encoding the +36 GFP described by McNaughton *et al* [17] fused to human calmodulin cloned into the NcoI and BamHI wherein the synthetic gene possesses a 6xHis tag after an initiating Met-Gly encoded by the NcoI site.

Cargo protein genes were based on the series of supercharged enhanced GFPs described by Thompson *et al* [16] fused to a canonical calmodulin binding site (KRRWKKNFIAVSAANRFKKISSSGAL) and HiBiT sequence (VSGWRLFKKISGGSG) cloned into NdeI and BamHI sites so that the N-terminal His tag is followed by GFP then CBS then HiBiT in constructs denoted “GCH” or CBS then GFP then HiBiT in constructs denoted “CGH.” Rearrangement of the order of moieties in the fusion was necessary for reasons described below. In no case were His tags removed. CBS and HiBiT sequences contributed to net protein charge. Proteins used in this study are described in Table 1.

**Table 1.**
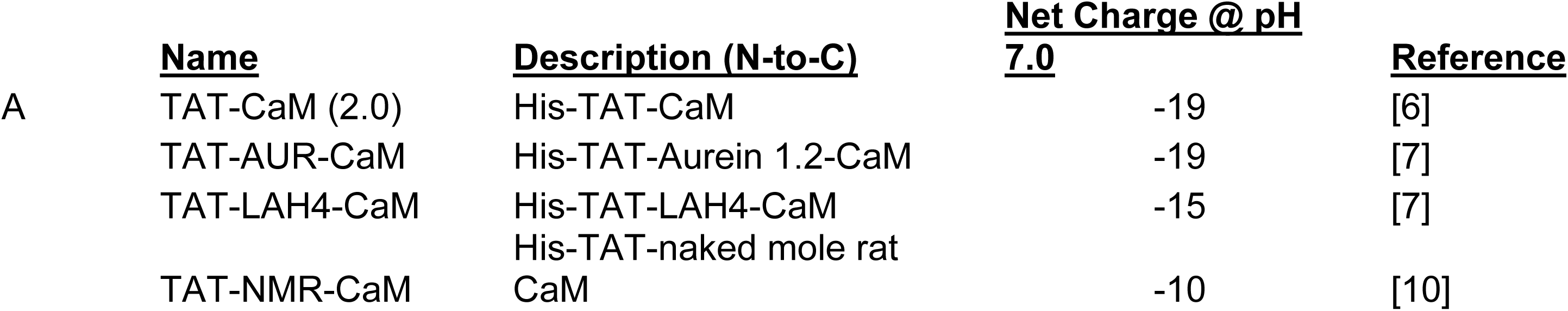
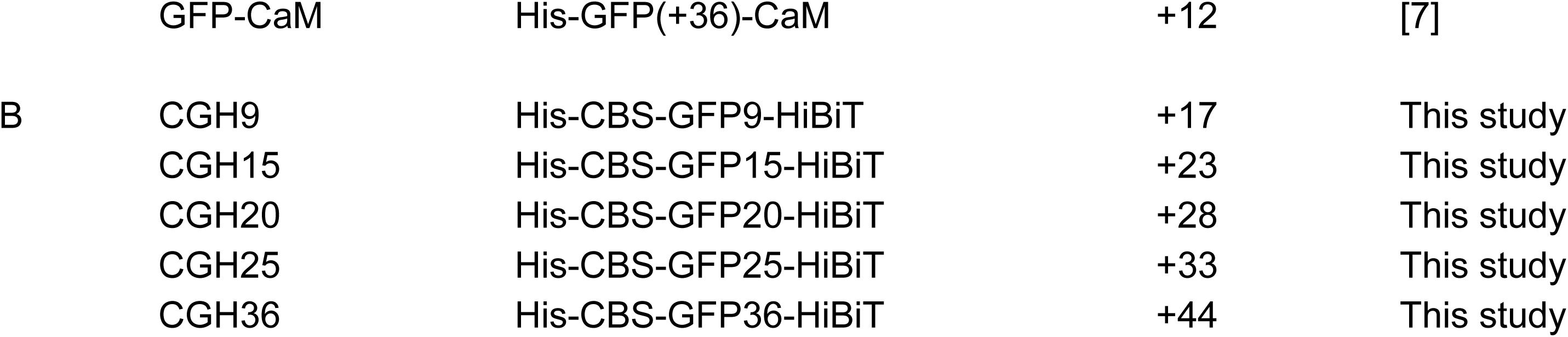
Important proteins used in this study. A, CPP- and CPPr-adaptors; B, cargo proteins. Sequences and additional information can be found in Supporting Information (S1 Fig).

### Expression, purification and labeling

Proteins were expressed essentially as described [5, 6] with minor modifications. Briefly, overnight cultures grown from single colonies were subcultured into 750 ml Luria-Bertani broth and grown with vigorous shaking at 37°C. At OD_600_ ∼0.4, temperature was lowered to 30°C and cells were induced with 0.2 mM IPTG and growth continued for four hours. Cells were harvested by centrifugation at 10,000 x g and frozen at -80°C.

Purification of CPP-adaptors was performed as described [5]. Supercharged GFP cargos and GFP-CaM required protocol modification due to the tendency of the highly positive GFP cargos to precipitate in low salt. Cell pellets were thawed on ice, resuspended in 2M NaCl lysis buffer (50 mM Tris pH 8, 2 M NaCl, 2.5 mM imidazole, 10% glycerol) with Halt Protease Inhibitor Cocktail (ThermoFisher) added to 1x per manufacturer’s protocol. Cells were broken via passage through a French press at 20,000 psi and subjected to centrifugation at ∼27,000 x g for 30 minutes at 4°C to pellet unbroken cells and debris. Clarified lysate was passed over a 1 or 5 ml Fast-flow HisTrap cobalt affinity column equilibrated with 1M NaCl Lysis Buffer (50 mM Tris pH 8, 1 M NaCl, 2.5 mM imidazole, 10% glycerol) using an FPLC system and washed with wash buffer (1 M NaCl lysis buffer, with 10 mM imidazole) until baseline absorbance was attained, after which protein was eluted with elution buffer (1 M NaCl lysis buffer with 250 mM imidazole). Cargo containing fractions were sampled for analysis then frozen in liquid nitrogen and stored at -80°C.

Although some GFP cargos were sufficiently pure for use after the HisTrap column, most cargos were further purified with Calmodulin Sepharose 4B matrix (Cytiva) by open phase affinity column purification. For this purification, IMAC elution fractions were quick thawed in water, placed on ice and brought to 1 mM CaCl_2_ and 2 M NaCl. These fractions were then loaded onto a 1 to 3 ml column equilibrated with 2 M NaCl lysis buffer with 1 mM CaCl_2_, washed with 5 column volumes of the same and eluted with 1 M NaCl lysis buffer containing 10 mM EDTA. GFP-containing fractions were sampled for purity analysis, frozen in liquid N_2_ and stored at -80°C. Fractions were later thawed, concentrated to a concentration above 100 µM on Amicon Ultra concentrators with a nominal 30 kDa molecular weight cutoff. In preparation for fluorescent labeling, concentrated cargos were desalted in 0.5 or 2 ml Zeba (Pierce) spin columns into 10 mM HEPES, pH 7.4, 1 M NaCl, 10% glycerol and 1 mM CaCl_2_. Desalted cargo protein was quantitated and treated with 0.4-0.6 mole of DyLight 650 NHS Esters (ThermoFisher) per mole of cargo. Dye removal columns (ThermoFisher) were then used to remove unreacted dye as recommended by the manufacturer. Protein concentration was estimated by Bradford and fluor incorporation estimated with the algorithm suggested by the manufacturer based on spectroscopic concentration analysis at 280 and 652 nm. This procedure produced incorporation efficiencies of about 0.2-0.3 dye molecules/cargo. Labeled cargos were aliquoted, frozen in liquid N_2_ and stored at -80°C until use. Core comparative experiments were all performed with a matched set of doubly purified cargos with incorporation of 0.17 to 0.23 fluors per cargo molecule.

### Cell Culture

Baby hamster kidney (BHK) cells were purchased from ATCC (#CCL-10). Cells were maintained in a 37°C, 5% CO_2_ environment in growth media consisting of DMEM, GlutaMax (containing +4.5g/L D-glucose and 1.9 mM Ca^2+^ with no sodium pyruvate) and 5% fetal bovine serum (FBS). Cells were replated following trypsinization in chamber slides (Ibidi) in the same media 20 to 24 hours before use in cell penetration assays.

### Cell Penetration Assays

Internalization assays were modified from prior assays [6, 22] to accommodate the high salt necessary to prevent cargo precipitation without producing large osmotic effects on cells. Complex assembly was performed at room temperature for the same reason.

Adaptors stored in a standard storage buffer (10 mM HEPES, pH 7.4, 150 mM NaCl, 10% glycerol, 1 mM CaCl₂) and GFP cargos (plus GFP-CaM) stored in high-salt buffer (10 mM HEPES, pH 7.4, 1 M NaCl, 10% glycerol, 1 mM CaCl₂) were thawed and diluted to appropriate concentrations in their respective storage buffers.

Complexes consisting of CGH cargos and adaptors were assembled at 100× concentration by mixing 1 volume of cargo with 0.5 volumes of adaptor in their respective storage buffers, yielding a final salt concentration of 716 mM NaCl. Complexes involving CGH cargo and GFP-CaM were prepared in the 1 M NaCl storage buffer, followed by dilution with 0.5 volumes of the standard storage buffer yielding the same salt concentration.

Following assembly, complexes were incubated for 20-30 minutes at room temperature and then diluted using Glutamax with 5% FBS media, also at room temperature. Rapid dilutions were obtained by squirting 400 to 1000 ul of media directly at the complex solution in the bottom of a microfuge tube followed by pipetting up and down several times. After 3-4 minutes at room temperature media containing complexes were microfuged at maximum velocity (>15,000 rcf) for 2 min to remove precipitates, briefly warmed in a 37°C bath (3-4 minutes) and transferred onto BHK cells from which growth media had just been removed. Cells were placed into an open-lid container in the incubator for 3 min and then, with lid closed, transferred to the confocal microscope. Care was taken to maintain conditions as close as possible to 37°C under 5% CO_2_ during transfer to an atmosphere-controlled chamber on the confocal microscope under the same conditions.

Confocal analysis was conducted using a Zeiss LSM 900 microscope, typically at 400x magnification with an image width of 160 µm. Imaging parameters were optimized to maximize image quality, including the use of low laser intensity and Airyscan-optimized confocal settings. All imaging within an experiment was performed with identical settings. GFP-labeled cargo exhibited strong binding to the slide surface. Thus, images were captured at the mid-cell plane to minimize surface-related signal artifacts and avoid image degradation caused by precipitates. Typically, 4-6 wells with varying conditions, e.g. varied cargos and adaptors, were imaged in a repetitive cycle of 10-20 minutes. Imaging was begun as soon as practically possible (∼8 min). However, the primary analytic period was from 40-70 minutes, which minimized the time differential between the first and last images.

Time-course movies comparing cargo internalization in the presence and absence of TAT-CaM were conducted concurrently. Complexes prepared as above were carried to the confocal microscope with cells prepositioned in 8 well slides and added to wells immediately after removal of growth serum. Autofocus to maintain image clarity was set as fast as possible and imaging begun about 5 min after complex addition.

In general, confocal images were adjusted for presentation by raising signal intensity of the highest panels to the top of the scale. Fluorescent signals were adjusted concordantly within each experiment although in some experiment conditions signal from GFP in each panel was adjusted to isolate yellow extracellular cargo from much redder intracellular cargo. Uniform background was also reduced in som panels. In all cases, representative images are shown from experiments repeated at least three times each producing multiple profiles from consecutive imaging cycles.

## Results

### Intrinsic internalization properties

As preparation for analysis of adaptor stimulated cargo internalization, it was necessary to demonstrate the intrinsic internalization properties of supercharged GFP cargos, initially designed as His-GFP-CBS-HiBiT fusion proteins. Although not used in the current study, the HiBiT sequence in association with a cytoplasmic fragment of luciferase produces light upon endosomal escape as part of a quantitative endosome escape assay. [23]. We also note that the supercharged GFPs have a tendency to precipitate, necessitating purification and storage in high salt as described in Materials & Methods.

As expected, GFP cargos with net positive charge have cell penetration abilities that increased as the positive charge of the GFP increased (Fig. 1A). As recently noted by the originating lab [16], supercharged GFP internalization does not appear to be robust. We recognized that these internalized GFPs never display perinuclear localization as observed for cargos that have attached fluorescent labels. We reasoned this might be due to loss of GFP fluorescence caused by lysosomal proteolytic cleavage, contrasting with slow loss of the incorporated label that would require very complete degradation. Unfortunately, the original His-GFP-CBS-HiBiT constructs proved too difficult to purify and label intact, necessitating a redesign to His-CBS-GFP-HiBiT (CGH) structure that is more stable. To compare the behavior of variously charged CGH cargos, a matched set of identically purified cargos was labelled with limiting amine-reactive 650 DyLight NHS-Ester that incorporated only 0.2 dyes per CGH cargo molecule (S2 Fig).

**Fig 1.**
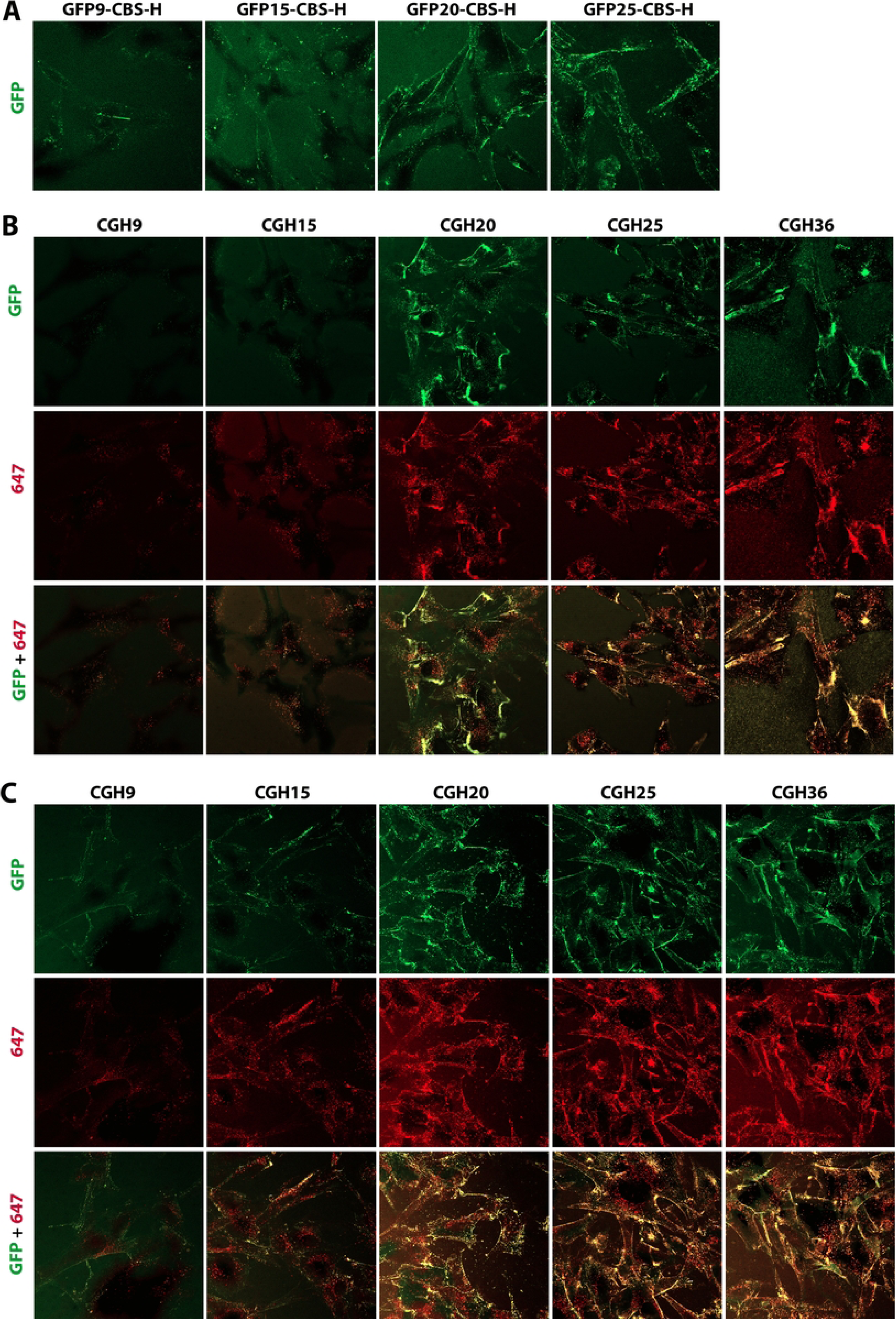
Intrinsic internalization of supercharged GFP cargos. Cargos were diluted with media and added to cells grown at 37^°^C under 5% CO_2_ for 22-26 h on 8-well coverslip slides. Slides maintained under these conditions were transferred to the confocal microscope and imaged continuously for up to 90 min in a compartment that maintained cells at 37^°^C under 5% CO_2_. **(A)** Internalization of supercharged GFP cargos with the design His-GFP-CBS-HiBiT and net GFP charge of +9, +15, +20 and +25. Representative profile (n=3) showing GFP internalization 40-70 minutes after addition of media containing 200 nM cargo to cells. **(B)** Representative profile (n=7) showing internalization of 100 nM far-red labeled GFP cargos with the design His-CBS-GFP-HiBiT (CGH) and net GFP charge of +9, +15, +20, +25 and +36 between 40-70 minutes. **(C)** Representative profile (n=6) showing internalization of far-red labeled CGH cargos at 400 nM cargo. A matched set of CGH cargos labeled with DyLight 650-NHS Ester at ∼0.2 moles dye per mole of cargo was used for direct comparison of 5 alternatively charged cargos. Image sets presented are representative of profiles from experiments with multiple profiles between 40-75 minutes. Confocal analysis was performed on a Zeiss 900 at 400x total magnification with simultaneous collection of GFP and far-red fluorescence for dual labeled cargos.

When the intrinsic internalization ability of these dual color CGH cargos was tested, the internalized GFP fluorescence pattern appeared similar to that observed with the original GFP constructs when one focuses on apparent endosomes (compare Fig.1A to Fig 1B, top panel). However, the far-red label on the cargo showed dramatically more intracellular localization (Fig 1B, middle panel), which also increased with increasing net GFP charge from +9 to +36. Adjusting the intensity of the green signal at the cell surface so that the false red signal is arbitrarily equal to the green signal segregates surface from most intracellular complexes. This is true even though these images were taken while cargos were present in the media. While pondering the extreme difference between GFP and false-red patterns, we realized that GFP fluorescence is pH-dependent and will be quenched below pH 6.2, as expected in most endosomal environments [24]. On the other hand, surface-localized complexes associated with the outer cell membrane (Fig. 1B) are in the neutral pH extracellular environment and should produce full GFP fluorescence. There are punctate structures that retain yellow coloration; however, in most cases these appear to represent membrane structures in contact with extracellular environment as they are found lining the cell edge or in disturbed membrane environments (Fig.1B). Thus, loss of most endosomal GFP fluorescence isolates red endosomal coloration, producing a readout that is extremely good evidence that cargo is present in acidic intracellular endosomes and not on the cell surface. Focusing only on red endosomes in the dual color panel (Fig.1B, bottom panel), it is apparent that CGH20 internalization is less than internalization of CGH25 or CGH36.

The experiment above (Fig. 1B) was performed at cargo concentrations of 100 nM, which is 10-fold or more below the concentrations typically used with covalently linked CPP-cargos. When the same supercharged CGH cargos were used to treat cells at 400 nM (Fig. 1C), the pattern of maximum internalization shifted so that less-charged CGH cargos (CGH15 and CGH20) displayed a relative increase in internalization. For these supercharged CGH cargos, internalization often correlates with cell surface labeling and both surface labeling and internalization appear saturable. This correlation suggests that charge increases internalization by increasing surface association. This makes sense as negatively charged proteoglycans on the cell surface are believed to be essential to the function of positively charged CPPs [25].

### INTERNALIZATION OF CGH CARGO SERIES WITH A SINGLE ADAPTOR

To demonstrate relative effects of adaptors on internalization of less positive cargos, internalization of the 5 supercharged CGH cargos were directly compared in the presence of 5 adaptors previously shown to be efficacious. Obviously, confocal images must be sequentially collected, creating an unavoidable time differential between the first and last images. Nevertheless, sequential sets create time series that show increasing internalization at longer cell treatment times. Such a time series for internalization of the CGH cargo series at 100 nM in the presence of excess of TAT-CaM is shown in Figure 2. In this experiment CGH25 was imaged first as soon as possible after complex addition (9 min) to show the low level of internalized CGH25 in the first image set relative to later sets. However, even in the set begun at 29 minutes, the image order is no longer obvious. Sets taken between 40-70 minutes are generally similar, suggesting the internalization process has reached a relatively stable profile pattern. This is only approximate, and slowly internalizing cargos do display a relative increase at longer times. To make the internalization data as simple and comparable as possible, we focus on the representative sets taken between 40 and 65 minutes. The time courses are included as supplemental figures both to support these profiles and because they often include additional information.

**Fig. 2.**
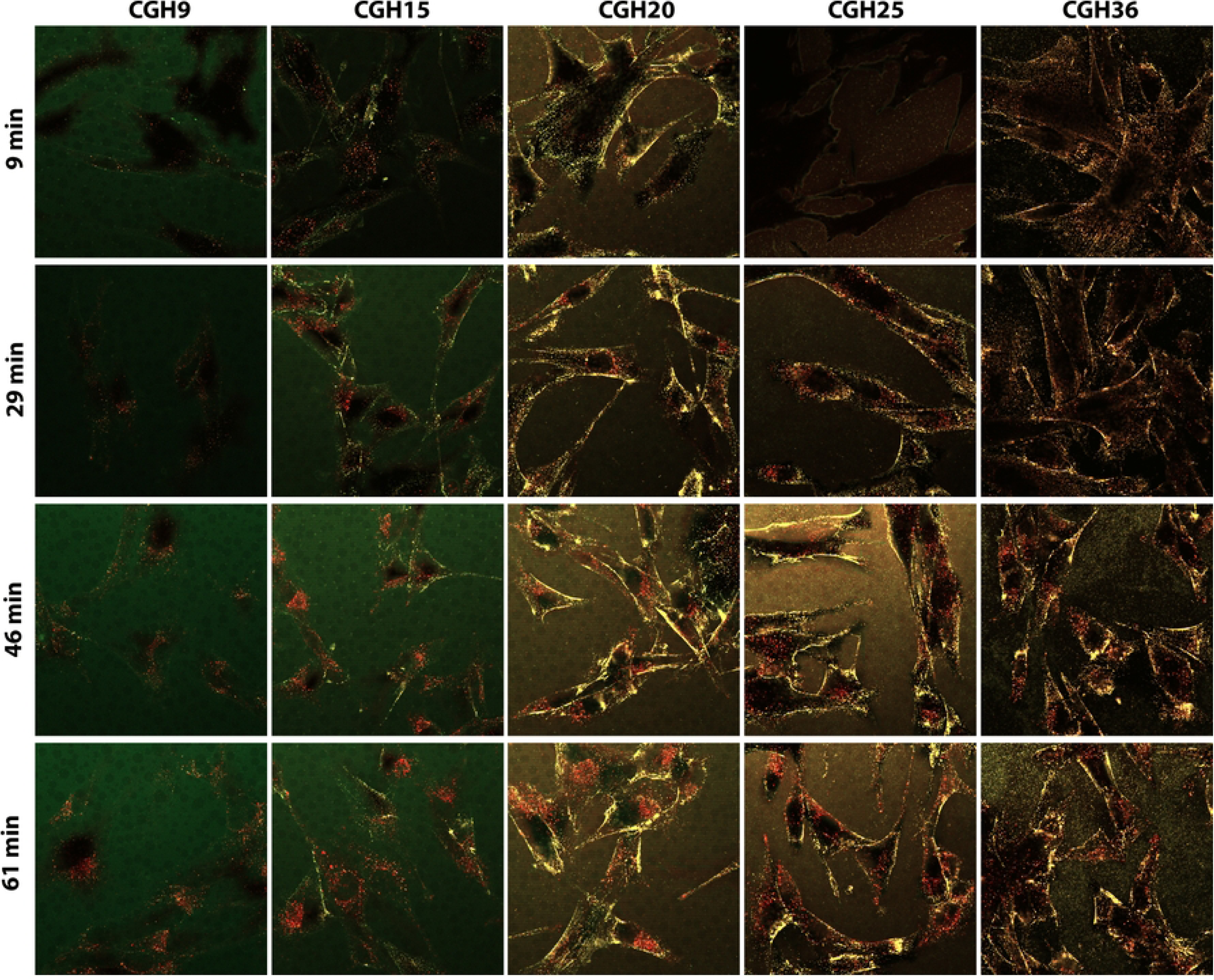
Time course of supercharged CGH cargo internalization in complex with TAT-CaM. Far-red labeled CGH cargos were combined with TAT-CaM as a premix (see Methods) so that dilution in media resulted in 100 nM cargo with 150 nM TAT-CaM. Adaptor/cargo complexes in media were handled as described in Methods, brought to 37°C and added to cells grown on coverslip slides at 37°C under 5% CO_2_ and then maintained under these conditions. To show the effect of time on cargo internalization, cells containing CGH25 complex were imaged first at 9 minutes after complex addition. Later sets were imaged beginning 29 min, 46 min and 61 min after complex addition with the cycle order of CGH25, CGH36, CGH9, CGH15, CGH20. For presentation in all images GFP is green and far-red 647 dye is false-red.

As shown above, the intrinsic internalization of supercharged cargos is both charge- and dose-dependent. A side-by-side comparison of TAT-CaM-mediated internalization of the 5 CGH cargos at 100 nM (Fig. 3A and S3A Fig) and 400 nM (Fig. 3B and S3B Fig) show profiles that match one another and are similar to the intrinsic internalization of the CGH series at 400 nM (Fig. 1C). Thus 100 nM cargos with TAT-CaM increased internalization of less charged CGHs relative to intrinsic levels, while 400 nM CGH cargos with TAT-CaM show little change compared to cargos alone. The parallel experiment with the adaptor TAT-AUR-CaM showed little adaptor effect as internalization profiles were similar to intrinsic internalization of the CGH series at both 100 nM (Fig. 3C and S3C Fig) and at 400 nM (Fig. 3D and S3D Fig). For both of these adaptors, failure to increase internalization of less charged cargos correlates with limited surface association. However, this relationship is not absolute and there is more variability in surface association than in internalization.

**Fig. 3.**
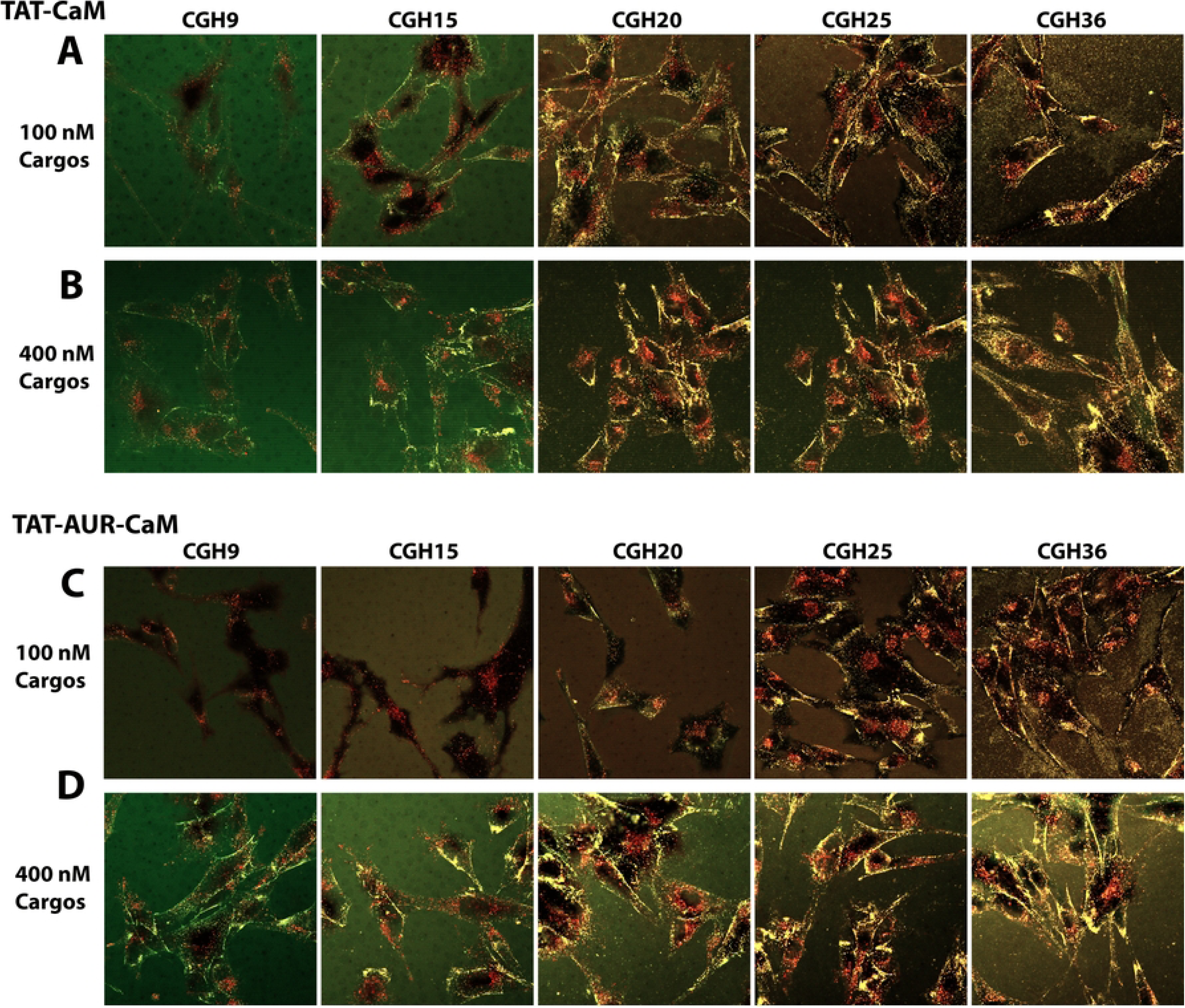
TAT-CaM and TAT-AUR-CaM do not drive maximal Internalization of all CGH cargos. CGH cargos and adaptors were combined in a premix and then diluted in media followed by addition to cells and incubation at 37°C under 5% CO_2_. For these experiments TAT-CaM was in 1.5-fold excess of CGH cargos at **(A)** 100 nM and at **(B)** 400 nM, while TAT-AUR-CaM was in 1.1 fold excess of CGH cargos at **(C)** 100 nM and **(D)** 400 nM. Image sets presented are representative of profiles from 3 experiments with multiple profiles between 40-75 minutes.

In contrast, for the adaptors TAT-LAH4-CaM, GFP-CaM and NMR-CaM, cargo internalization became much less sensitive to cargo charge. Most dramatically, TAT-LAH4-CaM internalized all CGH cargos at 100 nM (Fig. 4A and S4 A-B) and at 400 nM (Fig 4B and S4 A-B Fig), although the larger data set suggests CGH9 at 100 nM trended lower than other cargos dues to less rapid entry. Robust internalization was accompanied by similar surface loading of all TAT-LAH4-CaM/CGH complexes.

**Fig. 4.**
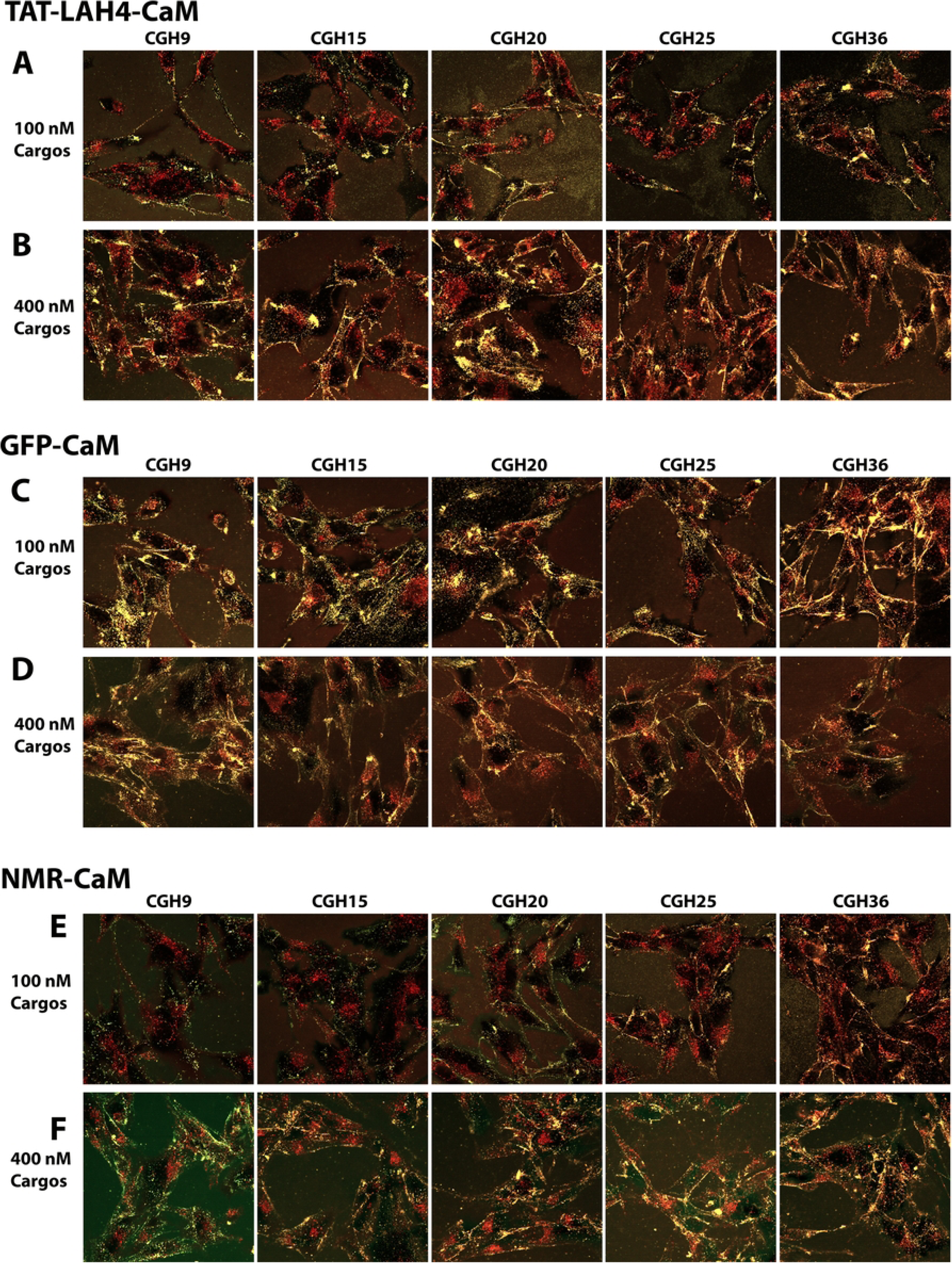
TAT-LAH4-CaM, TAT-NMR-CaM and GFP-CaM cause maximal internalization of less charged CGH cargos. CGH cargos and adaptors were combined in a premix and then diluted in media followed by addition to cells and incubation at 37°C under 5% CO_2_. For these experiments all adaptors were in excess of CGH cargos. Profiles using 1.5x TAT-CaM with CGH cargos at **(A)** 100 nM and at **(B)** 400 nM, 1.1x GFP-CaM with CGH cargos at **(C)** 100 nM and **(D)** 400 nM and 1.1x TAT-NMR-CaM with CGH cargos at **(E)** 100 nM and at **(F)** 400 nM. Image sets presented are representative of profiles from 3-4 experiments with multiple profiles between 40-75 minutes.

Complexes of all CGH cargos with GFP-CaM also showed increased surface association and charge independent internalization at both 100 nM (Fig 4C and S4 C-D Fig). and 400 nM (Fig. 4D and S4 C-D Fig). Complicating interpretation, the increase in green signal due to the presence of two GFP domains within each complex made it more difficult to isolate red endosomal populations from dual color yellow populations. Decreasing green signal intensity failed to change potential intracellular punctates from yellow to red and it is likely that many are surface associated. Focusing on the red endosomes suggests a trend toward lower CGH9 internalization at 100 nM (Fig. 4C).

The parallel experiment with the TAT-NMR-CaM adaptor showed charge independent internalization of all cargos at 400 nM (Fig. 4F and S4 E-F Fig.) accompanied by consistent if modest surface loading. For cargos at 100 nM, a modest reduction in the internalization of less charged cargos (Fig. 4E and S4 E-F Fig.) correlates with reduced surface association. For reasons that are unclear, variability in the internalization at 100 nM was high and the profile shown is a consensus choice. Despite the caveats, all three of these adaptors produced complexes where CGH internalization appeared to overcame less positive cargo charge, resulting in similar levels of internalization.

### Individual CGH cargos internalized with an adaptor series

The effects of individual adaptors on the panel of supercharged CGH cargos allowed relative comparison of each adaptor’s effects on a set of variably charged cargos. These data suggested a grouping of adaptors into those that poorly increase internalization of less supercharged cargos and those that appeared to stimulate internalization of these cargos. However, these images do not demonstrate the ability of an adaptor to specifically increase internalization of each cargo. This requires the intrinsic ability of each cargo to be directly compared to adaptor induced internalization. Because adaptor induced internalization will generally be hidden if the CGH cargos were already at saturating concentration, we tested specific internalization at 100 nM where most CGH cargos are below intrinsic saturation levels. The intrinsic internalization of each CGH cargo was compared to that of complexes with all 5 adaptors in moderate excess. The most representative profiles for each cargo from multiple experiments were selected from the period between 40-70 minutes and combined into a single image (Fig. 5A-E). Note that Figure 5, column 1 contains only cargo and shows intrinsic internalization at 100 nM.

**Fig. 5.**
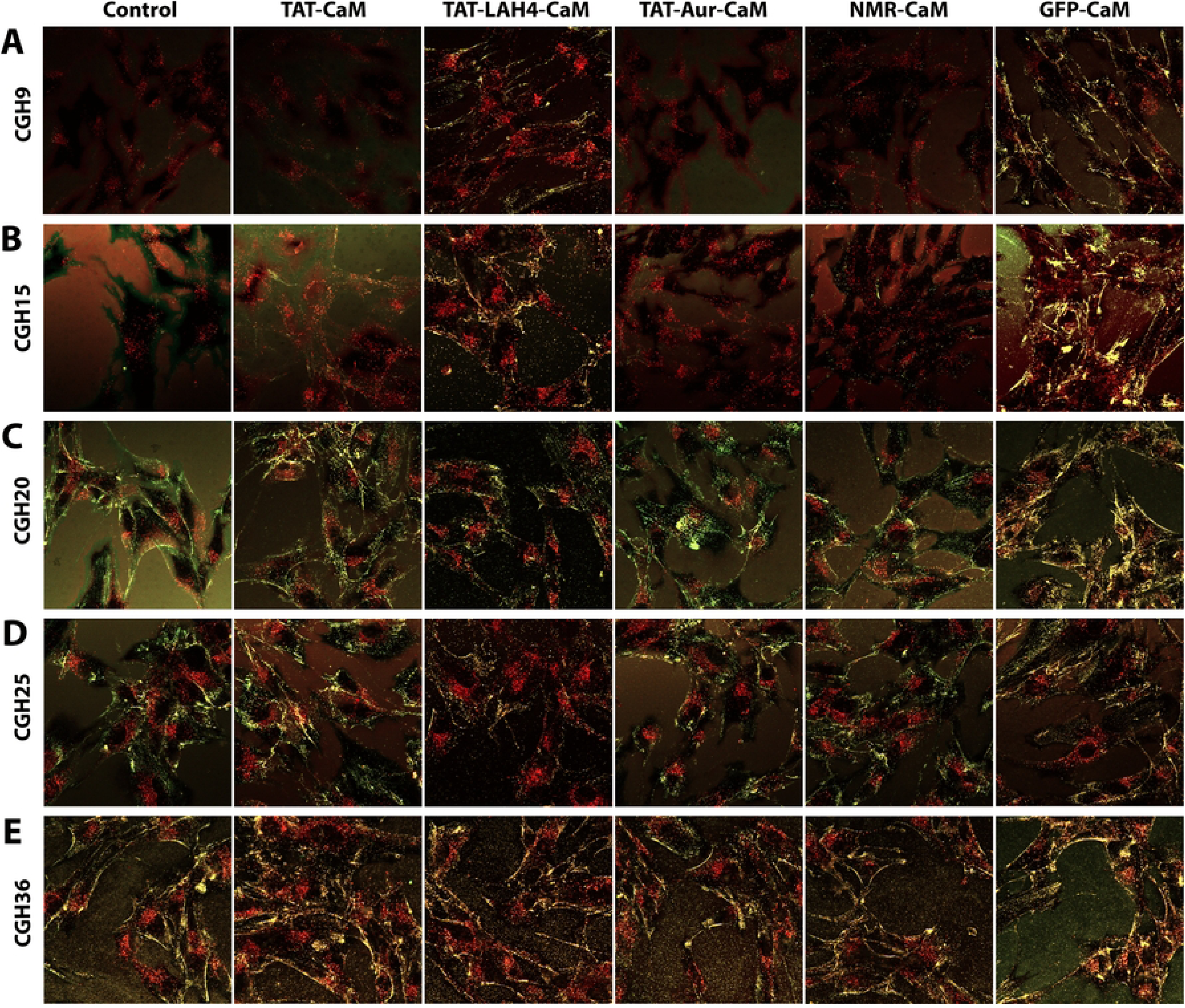
Comparison of intrinsic cargo internalization to that of cargo bound to TAT-CaM, TAT-LAH4-CaM, TAT-AUR-CaM, TAT-NMR-CaM and GFP-CaM. **(A)**CGH9 alone or combined with 5 adaptors in a premix was diluted in media to 100 nM with excess adaptor, followed by addition of the complex to cells and incubation at 37°C under 5% CO_2_ during imaging. Similarly, the cargos **(B)** CGH15, **(C)** CGH20, **(D)** CGH25 and **(E)** CGH36 either alone or combined with one of 5 adaptors in a premix was diluted in media to a concentration of 100 nM with excess adaptor. Image sets presented are representative of profiles from 3 experiments with multiple profiles between 40-75 minutes. Additional details provided with full experimental time course in the supplemental figures.

The most striking feature of this composite image is the robust internalization of most CGH cargos by TAT-LAH4-CaM (Fig. 5, column 2) including dramatic increases for CGH9 (Fig 5A and S5A Fig) and CGH 15 (Fig. 5B and S5B Fig) relative to intrinsic levels (Fig 5, compare column 3 to column 1). Internalization of CGH20 (Fig. 5C and S5C Fig) and CGH25 (Fig. 5D and S5D Fig) was also increased above intrinsic levels, while internalization of CGH36 (Fig 5E and S5E Fig) was similar.

In contrast, TAT-CaM actually inhibited CGH9 internalization (Fig. 5A, compare columns 1 and 2), consistent with the apparent lack of internalization observed above (Fig. 3A-B). Nevertheless, TAT-CaM consistently stimulates internalization of more positively charged cargos relative to their intrinsic internalization. Of note, TAT-CaM even produced a trend toward increased internalization of CGH36. TAT-AUR-CaM was similar in that it did not stimulate internalization of CGH9 but produced profiles similar to no adaptor controls for other cargos.

Although GFP-CaM appeared to stimulate internalization of less-positive CGH cargos in the profile above (Fig 4C), the data here indicate the situation is more complicated. While GFP-CaM does increase CGH9 (Fig. 5A) and CGH15 (Fig. 5B) internalization relative to intrinsic levels (compare columns 5 and 1), the adaptor decreases internalization of the highly charged CGH25 (Fig. 5D) and CGH36 (Fig. 5E) cargos below intrinsic levels. As a consequence of the divergent effects observed for 100 nM CGH cargos, GFP-36-CaM produced a fairly even distribution of intracellular cargos (Fig. 5 A-E, GFP-CaM in col 5) with CGH9 being modestly lower. As discussed above, the presence of two GFPs per complex necessitated additional reduction of the EGF signal to segregate extracellular (yellow) and intracellular (red) complexes.

While TAT-NMR-CaM did not stimulate internalization of CGH9 or CGH15, it decreased internalization of the most positive cargos, CGH25 and CGH36 (Fig 5, compare columns 5 and 1). This produced a less charge dependent profile as reported above (Fig. 4E). While inhibition of cargo internalization does not suggest use of these adaptors under these conditions, both TAT-NMR-CaM and GFP-CaM were specifically designed to increase cell surface association suggesting use at even lower concentration may be appropriate. Although not a useful topic of investigation, we have observed that most adaptor/cargo complexes produce inhibitory effects at too high a concentration.

Among the adaptors tested, the general effectiveness of TAT-LAH4-CaM with relatively neutral cargos at low concentrations is striking. However, it is the ability of TAT-CaM to increase internalization of positively charged CGHs that was most surprising. We previously showed that TAT-CaM internalizes near neutral cargos [7], but only at µM concentrations typically used with covalently linked CPPs. Further, TAT-CaM failed to increase internalization of positively charged proteins including Cas9 and tamavidin [7]. To explore the ability of TAT-CaM to internalize positive proteins at varied concentrations, we tested internalization of TAT-CaM/CGH20 complexes (Fig. 6B) compared to CGH20 alone (Fig. 6A) and found that TAT-CaM increased internalization at every concentration. This ability of TAT-CaM was unexpected as the utility of this adaptor had been limited to concentrations in the µM range; however, those experiments had been done using a near neutral cargo [7]. We also tested whether excess TAT-CaM beyond the cargo concentration impacted internalization and observed no effect from varied adaptor in moderate excess of CGH20 concentration (Fig. 6C).

**Fig. 6.**
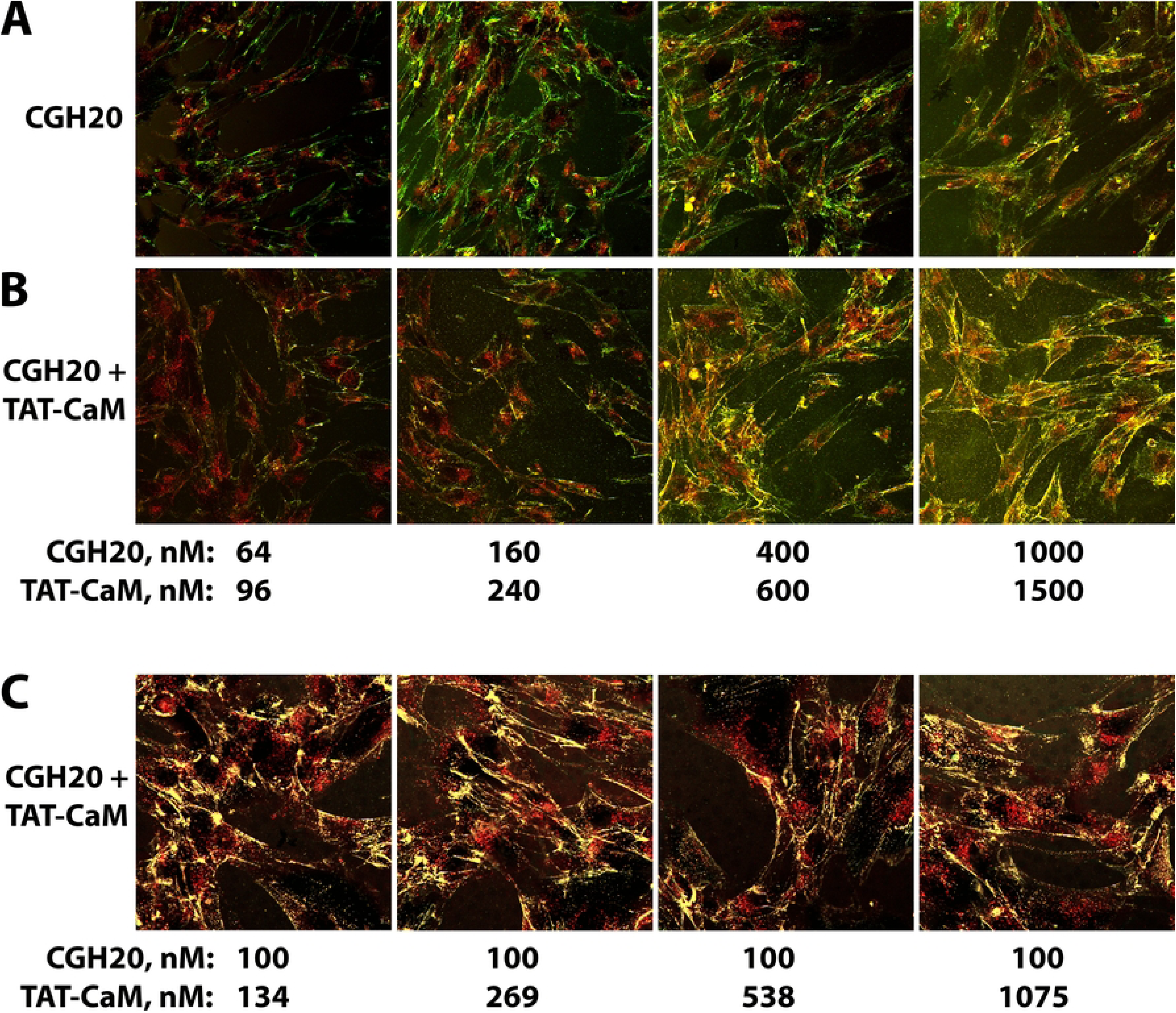
TAT-CaM increases internalization of CGH20 across a range of complex concentrations. CGH20 and TAT-CaM were combined in a premix and then diluted in media to the concentrations indicated, followed by addition to cells and incubation at 37°C under 5% CO_2_. Comparison of CGH20 internalization **(A)** without or **(B)** with TAT-CaM at a 1.5-fold excess of CGH20 cargos. Images ± TAT-CaM taken consecutively at each concentration. Image taken at 200x with image width of 320 µm. **(C)** Internalization of 100 nM CGH20 with 4 concentrations of excess TAT-CaM. Image sets presented are representative of profiles from 3 experiments with profiles between 40-75 minutes. Imaging order: high to low concentration.

Even though the cell penetration experiments above include a time course, results are still based on endpoints beyond 40 minutes of incubation. Although it is impractical to analyze confocal experiments using movies, limited validation of our endpoint assay was obtained for TAT-CaM complexes with the 5 CGH cargos by simultaneously imaging internalization with and without TAT-CaM. We chose the 400 nM cargo concentration to clearly demonstrate TAT-CaM stimulated early internalization above intrinsic velocities. Static images of selected time points in these experiments are shown in Figure 7 A-E and the movies are included as supplemental figures (Supp. Fig. 6A-E). As suggested by the static images, the movies for CGH15, CGH20 and CGH25 show TAT-CaM increased internalization above intrinsic levels with stimulation being more apparent early in the time course. As observed above (Fig. 5), CGH9 internalization appears to be inhibited by TAT-CaM, at least at longer times. Stimulation of CGH36 internalization by TAT-CaM is at best modest.

**Fig. 7.**
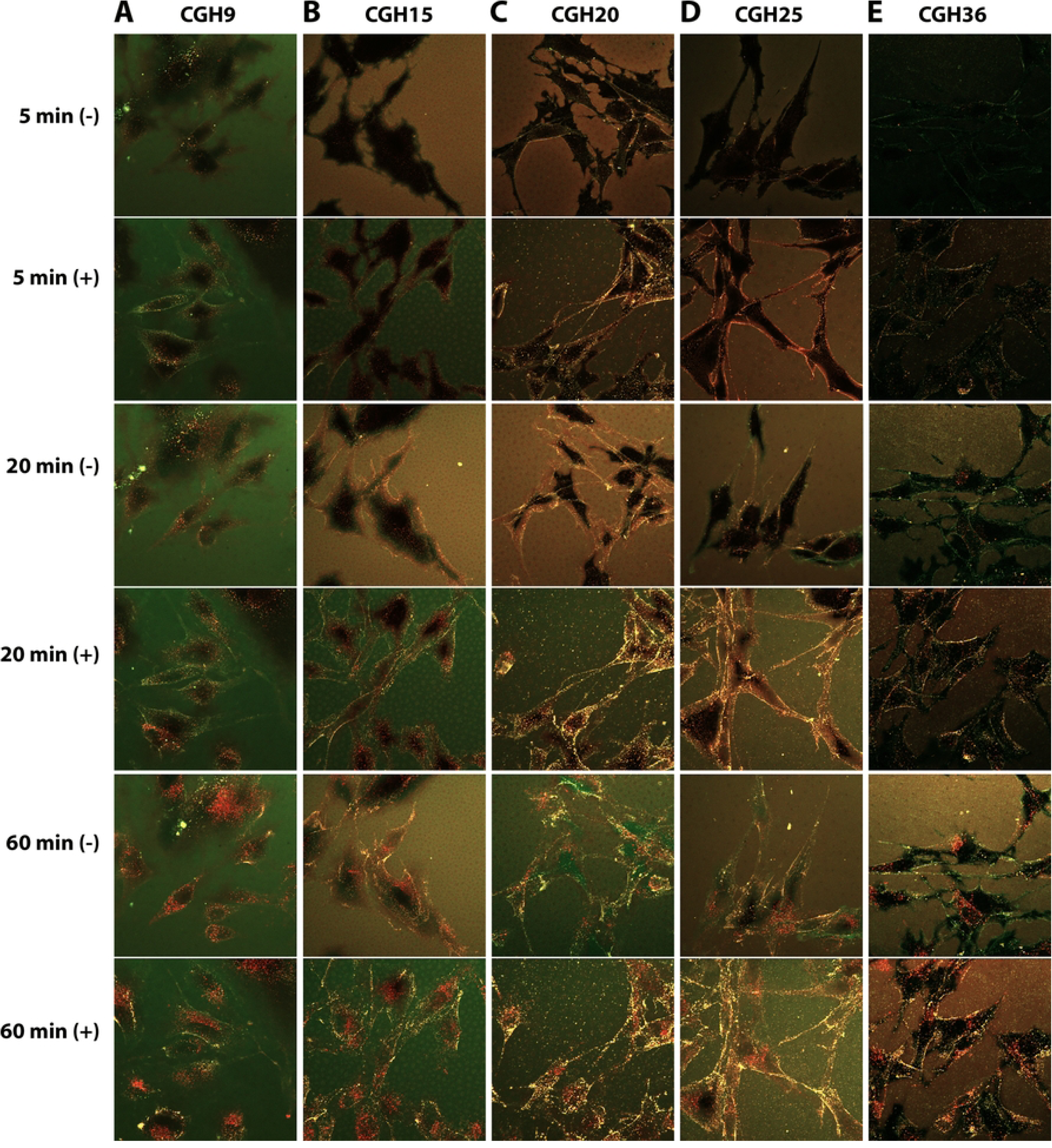
Selected still images from simultaneous movies of each cargo with or without TAT-CaM. For each cargo, movies showing internalization were initiated by addition of media with 400 nM CGH cargo without (-) or with (+) 600 nM TAT-CaM. These movies were obtained by alternative imaging of (-) and (+) wells for at least 60 minutes. Paired still images at selected times during the movies show internalization of **(A)** CGH9, **(B)** CGH15, **(C)** CGH20, **(D)** CGH25 and **(E)** CGH36, without or with TAT-CaM. Full movies are available as supplemental figures: S6A, S6B, S6C, S6D and S6E.

In contrast to TAT-CaM, the adaptor, TAT-LAH4-CaM, was effective at stimulating internalization of CGH cargos regardless of GFP charge state. Given the ability of TAT-CaM to stimulate CGH20 internalization beyond maximal intrinsic levels, we wished to determine if TAT-LAH4-CaM had this same ability. While TAT-LAH4-CaM stimulated CGH20 internalization most strongly at lower complex concentrations (compare Fig 8A to 8B), even at 800 nM CGH20, the complex specifically increased cargo internalization (compare Fig 8A to 8B, far right). As with TAT-CaM, TAT-LAH4-CaM in excess of cargo concentration had little effect on internalization of complexes with 100 nM CGH20 (Fig. 8C) or with 100 nM CGH15 (Fig. 8D).

**Fig. 8.**
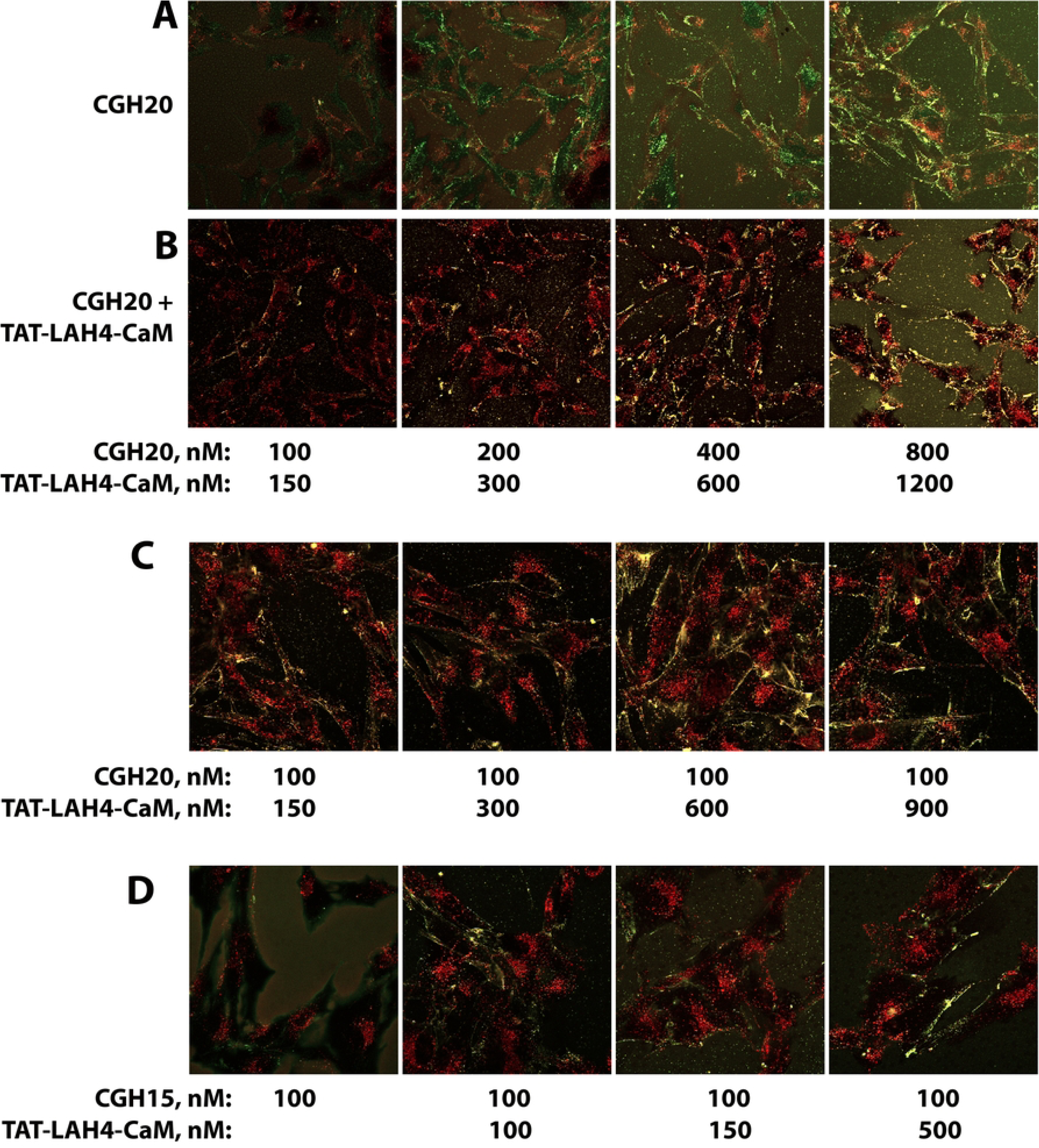
TAT-LAH4-CaM specifically increases internalization of CGH20 across a range of complex concentrations. CGH20 and TAT-LAH-CaM were combined in a premix and then diluted in media to the concentrations indicated, followed by addition to cells and imaging. Comparison of CGH20 internalization at 100, 200, 400 and 800 nM **(A)** without or **(B)** with a 1.5-fold molar excess TAT-LAH4-CaM. Images ± TAT-CaM taken consecutively at each concentration. **(C)** Internalization of 100 nM CGH20 with excess TAT-LAH4-CaM at 150, 300, 600 and 900 nM. **(D)** Internalization of 100 nM CGH15 without or with TAT-LAH4-CaM at 100, 150 and 500 nM. Image sets presented are representative of profiles from 3 experiments with profiles between 40-75 minutes. Imaging order: high to low concentration.

The adaptors TAT-NMR-CaM and GFP-CaM were designed to mitigate the negative charge of CaM and are very effective with some cargos [7, 10]. We suspected GFP-CaM/CGH complexes were saturating the cell surface to the point of inhibiting internalization and wanted to determine if reduced complex concentration would increase CGH internalization. The CGH15 cargo was used to avoid saturating effects from intrinsic internalization. As expected, across a range of very low concentrations (12.5nM-100nM) where this cargo alone internalizes poorly (Fig. 9A), GFP-CaM produced strong specific increases in CGH15 internalization (compare Fig. 9A to 9B). Unfortunately, at higher GFP-CaM/CGH15 complex concentrations, CGH15 internalization increased very little (Fig. 9D) while intrinsic CGH15 internalization increased dramatically (Fig. 9C) and exceeded that of the complex (Fig. 9C versus 9D). The similarity of GFP-CaM/CGH15 internalization from 25 nM to 400 nM suggests limits due to surface saturation combined with unknown characteristic of the GFP-CaM/CGH15 complex which prevents cargo internalization relative to CGH15 alone. The effect of 1.5 or 2 fold excess GFP-CaM above CGH15 concentration was also tested and did not appear to increase internalization (Morris, unpublished). Problematically, excess GFP-CaM both increased surface labelling and produced yellow punctates greatly complicating interpretation. The propensity of GFP-CaM to strongly associate with the plasma membrane suggests this adaptor in large excess would complete with GFP-CaM/CGH complexes.

**Fig. 9.**
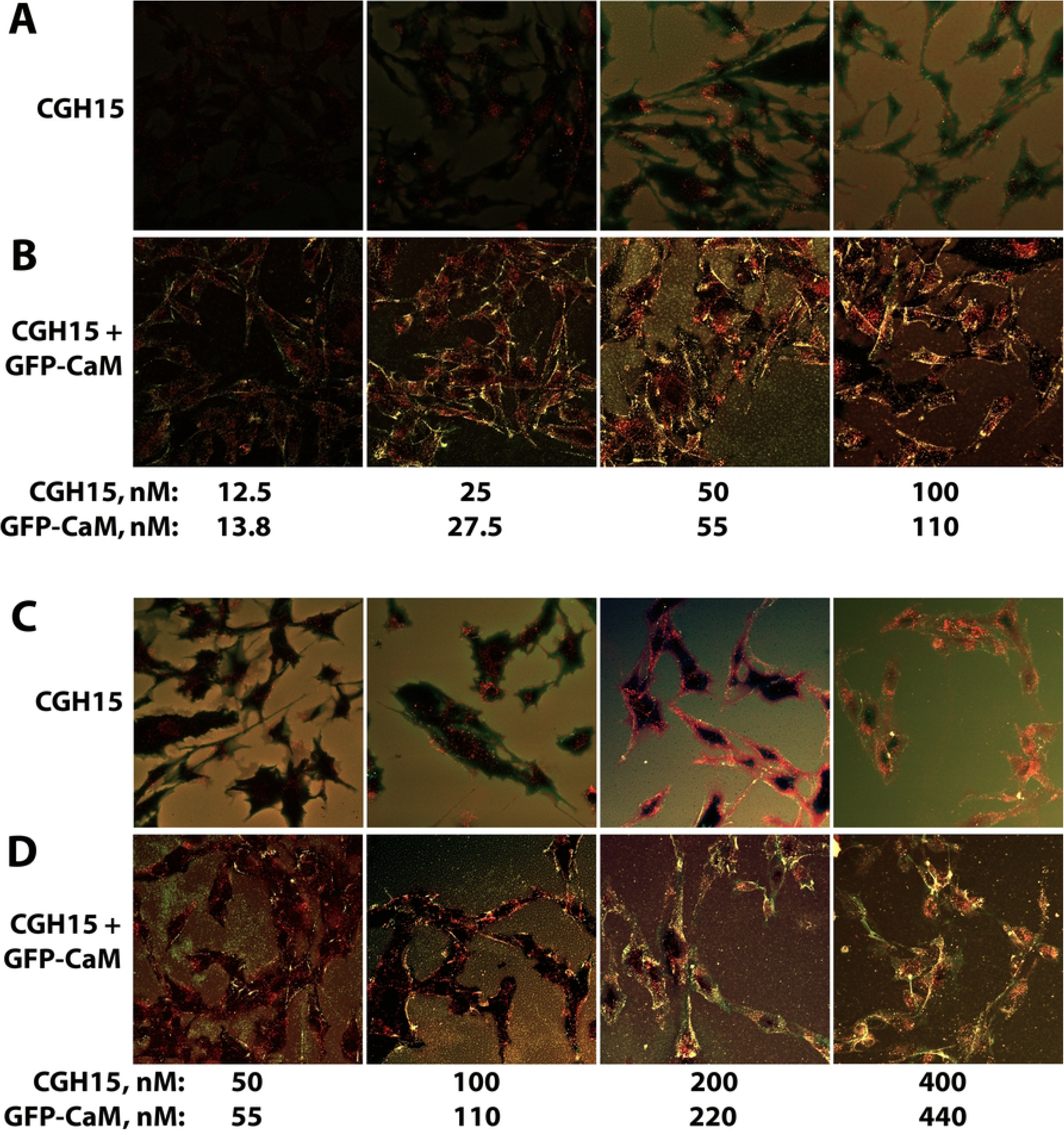
Dose dependence of CGH15/GFP-CaM internalization at low and high cargo concentrations. CGH15 and GFP-CaM were combined in a premix and then diluted in media to the concentrations indicated, followed by addition to cells and incubation at 37°C under 5% CO2. Low dose comparison of CGH15 internalization at 12.5, 25, 50 and 100 nM without **(A)** or with **(B)** a 1.1-fold excess of GFP-CaM. High dose comparison of CGH internalization at 50, 100, 200 and 400 nM without **(C)** or with **(D)** a 1.1-fold excess of GFP-CaM. Imaging order: high to low concentration with (-) GFP-CaM control taken immediately prior to CGH15/GFP-CaM complex.

## Discussion

The CPP-adaptor system was developed to solve the problem of endosomal entrapment by decoupling cargos from CPPs, which remain bound to endosomal membranes and trafficked to lysosomes. However, upon application it became clear intracellular separation of cargo from adaptor had numerous other advantages. Establishing the ability of an adaptor to increase specific internalization is a core part of our internalization assay and this feature soon revealed that cargos with highly positive net charge displayed intrinsic internalization. A particularly dramatic example was provided by the very positive cargo, Cas9, the protein component of CRISPR. Labeled TAT-CaM and labeled CBS-Cas9 internalized readily at concentrations more than 10-fold below TAT-CaM alone [7]. In other words, cargo mediated adaptor internalization.

Obviously, this challenged interpretations of many CPP internalization studies as the role of cargo is rarely tested in CPP studies. Our data showing that increasingly positive cargo charge increased internalization was strangely consistent, given that context specific effects must occur. This pointed to the need for systematic investigation of the effect of positive cargo charge and the interaction of cargo charge with CPP- and CPP-adaptor-mediated internalization.

### Intrinsic internalization

To address the effect of cargo charge on internalization, we designed cargos based on a set of mutated GFPs which had already been shown to display increasing internalization with increasing positive charge [16]. As reported, we found increasing GFP charge from +9 to +25 increased internalization; however, internalization appeared to be poor. When more stable redesigned CGH cargos were labeled with a covalently attached fluor, the label on these cargos revealed strong internalization despite a near absence of intracellular GFP fluorescence. The failure of intracellular GFP to fluoresce was almost certainly due to the acidic environment of intracellular endosome, which are generally below pH 6, where GFP fluoresces poorly.

While increased signal was valuable, the ability to distinguish dually fluorescing extracellular cargo from intracellular cargo that lacked strong GFP fluorescence was even more valuable. Confocal imaging could then distinguish yellow cargo present on the slide and outside the cell surface from red cargo inside the cell, removing the need for washing and the associated multilayered artifacts. As seen in our data, this segregation is not perfect, but in most situations it appears to be adequate.

### Experimental issues

While the dual fluor CGH cargos provided a powerful system for investigation of CPP mediated internalization, there are complications that seem to be intrinsic to the supercharged GFPs. Most significantly, cargos with highly charged GFPs tend to precipitate. Limiting precipitation required high concentrations of NaCl from the beginning of purification through dilution of adaptor/cargo complexes in media. High concentrations of cargos were then required so that large dilutions could limit salt addition to cell medium. Even so, some cargos and some complexes produced problematic levels of precipitation during imaging. Temperature also impacted internalization and experiments here were performed with minimal cooling of cells during handling and treatment. Given the difficulty of individual experiments and the complexity of the experimental space, systematic approaches were limited to variables that seemed particularly important. It is fortuitous that excess adaptor had little effect on complex internalization across a range of concentrations, removing one problematic variable.

Because confocal microscopes cannot simultaneously analyze multiple wells, a pseudo end point between 40 and 70 minutes was used for most analysis, despite the information lost regarding early internalization. Initial work with the CGH cargos revealed a charge dependent saturable surface association that usually correlated with maximal internalization. *A priori*, this suggests a limit on surface association due to positive charge and thus a limit on charge dependent internalization. Concentration was also an important determinant of surface loading and thus CGH cargo internalization. The relatively stable internalization profile observed across the CGH states between 40 and 70 minutes also appears to involve exhaustion of a dominant internalization process. The complex mechanisms that produce relatively stable profiles is beyond the scope of the current study.

### Adaptors

Given the dominant role of net cargo charge on intrinsic CGH internalization, the diversity of responses cargo charge produced by a limited set of 5 adaptors suggests that cargo charge is not the only determinant controlling internalization. In particular, TAT-CaM specifically increased internalization of most CGH cargos without increasing cargo localization at the cell surface. Indeed, cargo surface association appears to have trended somewhat lower with TAT-CaM relative to that observed with cargo alone. In addition, TAT-CaM was ineffective at increasing either surface localization or internalization of CGH9 and the limited increases in CGH15 internalization occurred with little surface localization. Endpoint assays showed that TAT-CaM was particularly effective with CGH20, resulting in strong internalization that appears to reach a common maximal level for the more positive cargos. Specific stimulation of CGH25 and CGH36 internalization was difficult to assess using the endpoint assay. Movies of internalization with and without adaptor showed that TAT-CaM specifically increased internalization of CGH25 and perhaps CGH36, although this was less obvious.

The ability of TAT-CaM to increase internalization of more moderately positive cargos such as CGH20 and CGH25 was striking. Further, the stimulatory effect of TAT-CaM on internalization occurred across a range of complex concentrations including some far below the µM range typically used with this adaptor. In contrast, TAT-CaM was very ineffective with the cargos CGH9 and CGH15, which displayed very little intrinsic internalization or membrane association. Consistent with the inability of TAT-CaM to increase surface association, a parsimonious explanation for this charge-dependent difference is that TAT-CaM stimulates internalization of cargos associating with the plasma membrane but has limited ability to recruit cargos to the cell surface.

In dramatic contrast, TAT-LAH4-CaM was impressive in driving maximal internalization of all cargos regardless of charge even at very low complex concentration. This ability correlates with a strong phenotype involving recruitment of CGH cargos of all charge states to the plasma membrane. While the LAH4 sequence was originally identified as an EP, later evidence revealed CPP activity [19, 26]. The TAT-LAH4-CaM adaptor was identified while investigating endolytic peptide (EP) functionality within the context of inclusion in both the adaptor (TAT-EP-CaM) and a maltose binding protein cargo (CBS-EP-MBP)[7]. As TAT-LAH4-CaM was able to internalize negatively charged MBP cargos it is not too surprising that TAT-LAH4-CaM increased surface association and internalization of neutral CGH15 and CGH9. Further, the TAT sequence in TAT-LAH4-CaM may be activating internalization as the adaptor stimulated CGH20 internalization even at high concentrations. TAT-AUR-CaM was developed in the same study and included here because the Aurein 1.2 sequence is endolytic [27] and because a CBS-AUR-MBP cargo produced a strong negative effect on GFP-CaM internalization associated with peripheral endosomal trapping [7]. While TAT-AUR-CaM may have had some inhibitory effects on CGH cargo internalization, the behavior was too modest to justify further study.

Both TAT-NMR-CaM and GFP-CaM represented successful efforts to improve adaptor function by increasing the net positive charge of adaptors [7, 10]. The dominance of charge and concentration in producing maximal intrinsic internalization of CGH cargos correlates with surface loading of a cargo. Consistent with this idea, GFP-CaM localized all of the CGH cargos to the cell surface and internalized these cargos to about the same extent at both 100 and 400 nM. Unfortunately, this equalization had as much to do with inhibition of internalization for more charged CGH cargos as it had to do with modestly increased internalization observed with CGH9 and CGH15. The idea that adaptor/cargo complexes can inhibit internalization at high concentration has been supported by our data for some time. Related to this, we have observed situations where surface bound cargo did not appear to internalize until washing and we have reported one instance of this involving GFP-CaM [7]. Analysis of the concentration dependence of GFP-CaM/CGH15 on complex internalization revealed strikingly little dose effect from 25 to 400 nM and combined with strong adaptor specific internalization at the low concentrations. Dose independent CGH15 internalization is consistent with strong surface association establishing maximal internalization and inconsistent with high dose inhibition. Unfortunately, maximal GFP-CaM-induced internalization is well below the intrinsic internalization of CGH15 at the highest concentrations, revealing some aspect of the GFP-CaM/CGH15 complex is problematic for internalization. Nevertheless, it is probable that adaptors with strong surface association can produce internalization at low concentrations similar to the maximal level achievable by CGH cargos themselves. While it would not seem to be necessary, robust internalization may require a trigger that stimulates the internalization process. Active investigation continues in this direction.

## Conclusion

Application of CPP-adaptor technology to a set of differentially charged cargos has produced a diverse set of outcomes that suggest the power this approach may deliver. At its core, the power of CPP-adaptors lies in the flexibility inherent to separation of the CPP function from the cargo, which allows different functionalities to be placed within the CPP and the cargo. We have uncovered an apparent limit to the maximal internalization that can be achieved by positive charge alone and shown that adaptors including TAT-CaM and TAT-LAH4-CaM may increase internalization to a higher level. We have also shown adaptors with high affinity for the cell surface can dramatically reduce the concentrations at which cargos will internalize. The cargos used in this study were designed with the HiBiT sequence that produces functional luciferase when HiBiT-containing cargo escapes the endosomes and combines with the remainder of the luciferase protein in the cytoplasm and that will be the next step. Those studies were delayed by the need to maximize CPP-adaptor internalization so that endosomal escape assays are not at the border of detection. In our recent studies, we have barely scratched the surface possible with CPP-adaptors using limited sets of cargos and adaptors. The development of effective CPP-mediated cargo delivery strategies will require optimization of internalization and endosomal escape. The next endeavor with the cargos and adaptors described in this study will be to measure and optimize parameters that enhance endosomal escape.

